# The Two-Component Nuclease-Active KELShedu System Confers Broad Anti-Phage Activity via Abortive Infection

**DOI:** 10.1101/2025.09.13.674623

**Authors:** Hengwei Zhang, Jiajia You, Hanwen Zhou, Zan Zhang, Hongxuan Wu, Di Zhang, Xuewei Pan, Weiguo Zhang, Xian Zhang, Zhiming Rao

**Author notes:** These authors contributed equally to this work (Hengwei Zhang, Jiajia You). Correspondence (X.Z.); (Z.R.).

## Abstract

Bacteriophages and bacteria engage in a continuous evolutionary arms race, driving the development of intricate bacterial defense systems such as CRISPR-Cas, BREX, Gabija, and Shedu. Here, we characterize a two-component KELShedu system in *Escherichia coli* that confers resistance to phages via abortive infection. The KELShedu system comprises KELA, a dsDNA-binding protein, and KELB, a metal ion-dependent nuclease harboring the DUF4263 domain. In addition, we find that physiological levels of NTP inhibit the DNA cleavage activity of the KELShedu system, suggesting that KELShedu’s activation depends on reduced intracellular NTP levels during phage invasion. Our research demonstrates that the KELShedu system responds to nucleotide depletion triggered by phage replication, leading to non-specific degradation of cellular DNA and ultimately inducing abortive infection. These insights into the KELShedu system expand the repertoire of bacterial anti-phage mechanisms and lay the groundwork for novel applications in microbial engineering and therapeutic development.

**Teaser:** The KELShedu system exerts anti-phage function by aborting infection in response to physiological changes following phage infection

## Introduction

Bacteriophages, viruses that infect bacteria, and their bacterial hosts are engaged in a constant arms race, driving the evolution of sophisticated defense and counter-defense mechanisms. Among these, systems such as CRISPR-Cas(*1, 2*), Restriction-Modification (R-M)(*3*), and the more recently characterized BREX(*4, 5*), Gabija(*6, 7*), and CBASS systems(*8*) play crucial roles in bacterial immunity. These systems employ a variety of strategies to detect and neutralize invading phages, often involving nuclease activities that degrade phage nucleic acids.

The initial step in any anti-phage defense is the detection of the invading phage(*9*). Various systems have evolved intricate methods to sense phage. The CRISPR-Cas system, a revolutionary discovery in molecular biology, provides adaptive immunity by capturing snippets of phage DNA and using them to recognize and cleave the phage DNA during subsequent infections(*1, 10*). The BREX (Bacteriophage Exclusion) system uses a combination of DNA methylation and restriction enzymes to recognize and degrade foreign DNA(*4, 11*). Central to this system is BrxR(*12, 13*), a protein that functions as a sensor and regulator, identifying phage DNA by its lack of specific methylation patterns. Similarly, the CBASS (Cyclic oligonucleotide-based Anti-phage Signaling System) employs signaling proteins that detect phage infection and activate downstream defense mechanisms(*8, 14*).

Once phage DNA is detected, bacterial cells deploy various effectors to neutralize the threat. In the BREX system, the effector component includes nucleases that degrade phage DNA, preventing the replication of the invader(*11*). Gabija, another defense system, employs a RecB-like nuclease that similarly targets and degrades phage DNA(*6, 7*), effectively halting the phage life cycle. The CBASS system activates upon phage infection, producing cyclic oligonucleotides that act as second messengers to trigger a range of antiviral responses, including the activation of nucleases(*8*) or membrane disruption (*14*). The nuclease activity is a common and crucial component of these systems, providing a direct means of neutralizing phage DNA.

In this study, we report the discovery and characterization of the KELShedu system, a novel two-component anti-phage mechanism that protects bacteria from aborting infection. This mechanism of abortion infection is different from that reported by Yajie Gu et al(*15*) on *Bacillus cereus* Shedu. We provide a comprehensive characterization of the KELShedu system, consisting of KELA and KELB proteins, detailing validating its function through a series of phage plaque assays, bacterial growth curve measurements, protein interaction assays and protein activity verification. We also found a mode of activation of the KELShedu antiphage system. After phage infection, the decrease of NTP caused by physiological activities such as phage replication and protein expression activate the DNA degradation activity of KELShedu system, which degrades intracellular DNA unspecifically and eventually leads to aborting infection. Our findings not only expand the repertoire of known anti-phage mechanisms but also lay the groundwork for future applications in biotechnology and synthetic biology to protect valuable bacterial strains from phage predation. The elucidation of the KELShedu system’s mechanisms contributes to the broader understanding of bacterial defense strategies and offers potential for developing new tools in microbial engineering and therapy.

## Results

### KELShedu is a two-component anti-phage system

Based on previous research, Shany Doron et al. reported that the Shedu system carries the core DUF4263 domain from *B. cereus* B4264, conferring protection against several *Siphoviridae* and *Podoviridae* bacteriophages, including phi105, rho14, spp1, and phi29(*16*). Building on this, we aimed to identify effective resistance systems that could help the industrial chassis cell *E. coli* BL21(DE3) resist bacteriophage infection(*17*). We selected proteins with the DUF4263 domain from various Enterobacteriaceae sources(*16*). We identified four Shedu proteins: KEL80639.1, BAI39103.1, UKB45259.1, and EHX83949.1 (Fig. S1). Notably, except for KEL80639.1 (KELB, 38.6 kDa), which forms a bicistronic operon with KEL80640.1 (KELA, 12.6 kDa), the other Shedu systems exist as monocistronic units. Therefore, we hypothesized that both KELA and KELB are essential for phage resistance and co-synthesized these genes. The codon optimization method for the KEL80639.1-KEL80640.1 operon (KELAB) is described in the Construction, Expression, and Purification section (Fig. 1A). We performed EOP assays with eight representative bacteriophages across four Shedu systems. Ultimately, we found that KELAB and BAI39103.1 conferred resistance to specific phages. KELAB exhibited significant resistance to T1, T7, and JNUWH1 phages (Fig. 1B, C), while BAI39103.1’s resistance was consistent with Wang et al.’s findings(*18*), showing resistance to T4, P251, and T7 phages (Fig. 1B).

**Figure 1.**
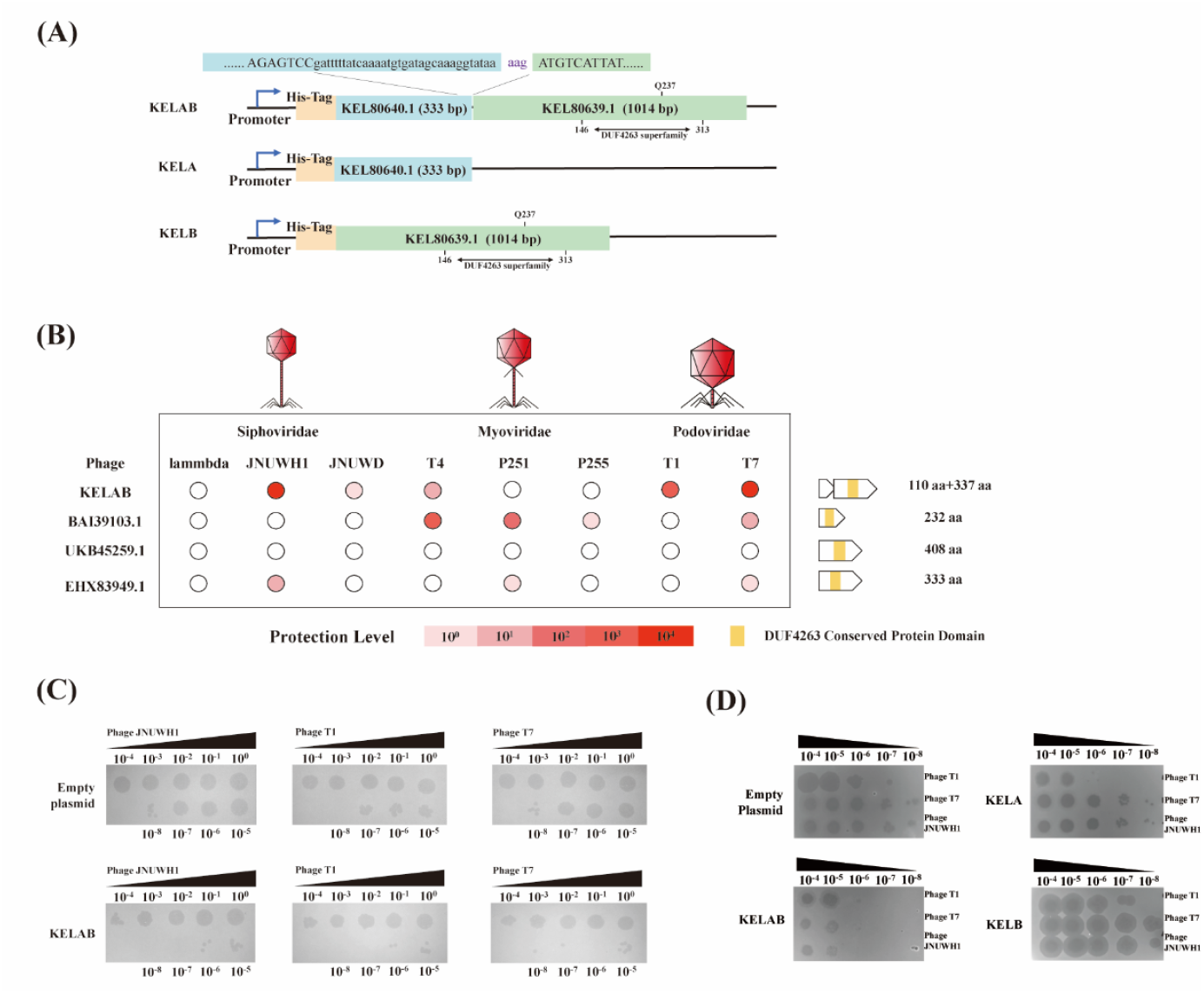
The two-component shedu system confers bacteriophage resistance to *E. coli*. (A) The KELAB gene cassette encompasses two genes, SduA (KELA, 333 bp) and SduB (KELB, 1086 bp), separated by a 3 bp intergenic region. The entire KELAB gene cassette, excluding the lowercase letters, has been codon-optimized. The DUF4263 superfamily domain is located within the 146-313 amino acid region of SduB. (B) Experimental validation of the shedu defense system from various sources. The protective efficacy was assessed using a serial dilution plaque assay, comparing strains harboring the shedu system with those containing an empty vector as a control. Data represent the mean of three independent experiments. Phage species are indicated above the corresponding names. To the right, the gene composition of the defense system is depicted, with the identified DUF4263 domain highlighted in yellow. Gene sizes are drawn to scale. (C) Expression of the KELAB cassette in *E. coli* BL21(DE3) confers resistance to phage infection. Shown are plaque assays with tenfold serial dilutions of phages T1, T7, and JNUWH1. The images are representative of three independent replicates. (D) Phage resistance in *E. coli* BL21(DE3) expressing different ORFs of the KELAB cassette. Strains containing KELA, KELB, the entire KELAB cassette, and empty vector were tested. Displayed are the results of plaque assays with tenfold serial dilutions of phages T1, T7, and JNUWH1. The images are representative of three independent replicates.

Given the unique nature of the KELAB operon, we validated the essential roles of both KELA and KELB in phage resistance. We separately expressed KELA and KELB in *E. coli* BL21(DE3) and evaluated their effects on resistance to phages T1, T7, and JNUWH1. As anticipated, the individual expression of either KELA or KELB within the KELShedu system significantly diminished the system’s broad-spectrum anti-phage activity. Interestingly, the expression of KELA alone conferred partial resistance specifically against T1 phage (Fig. 1D), potentially attributable to unique aspects of T1 phage entry mechanisms. These findings support that both KELA and KELB are indispensable for the anti-phage activity conferred by the Shedu system.

### KELShedu system mediates immunity through abortive infection

Abortive infection is a common bacterial defense mechanism against bacteriophages(*19*). Infected bacteria undergo premature cell death, disrupting the phage lifecycle and protecting the bacterial population. We investigated this mechanism by measuring the growth curves of *E. coli* expressing the KELShedu system compared to a control group lacking the KELAB genes. The assays were conducted at multiplicities of infection (MOI) of 0.1, 1, 10, and 100. The results demonstrated that in the control *E. coli* without KELAB, bacterial cultures collapsed within 20-60 minutes post-infection at MOI > 1 (Fig. 2A-F). In contrast, the KELShedu system conferred protection to *E. coli* at MOI 0.1 and 1, maintaining normal growth (Fig. 2A-F). However, at MOI 10, bacterial growth was arrested, and no further growth or lysis was observed, indicating that the KELShedu system effectively inhibited phage life cycle but also halted bacterial proliferation(*20*). At MOI 100, the phage-induced collapse occurred similarly in both KELAB-expressing and control cultures (Fig. 2BDF), suggesting a threshold limit for KELAB-mediated resistance, consistent with plaque formation observed in high phage concentration EOP assays. Although bacteria carrying the KELShedu defense system collapsed at an MOI of 100, similar to the control group, we quantified the phage titers in the resulting lysates under the same conditions. We observed that the phage titer in the lysates of KELShedu-expressing bacteria was at least two orders of magnitude lower than that in the lysates of bacteria containing an empty plasmid (Fig. 2G). To further rule out the possibility that the observed difference in phage titer was due to variations in phage adsorption rates, we compared the adsorption rates of phages to *E. coli* with and without KELAB. No significant difference was observed, indicating that KELAB allows phage attachment but protects bacteria through downstream interference post-attachment (Fig. S2). This result suggests that KELShedu aids in bacterial resistance against phages through an abortive infection mechanism.

**Figure 2.**
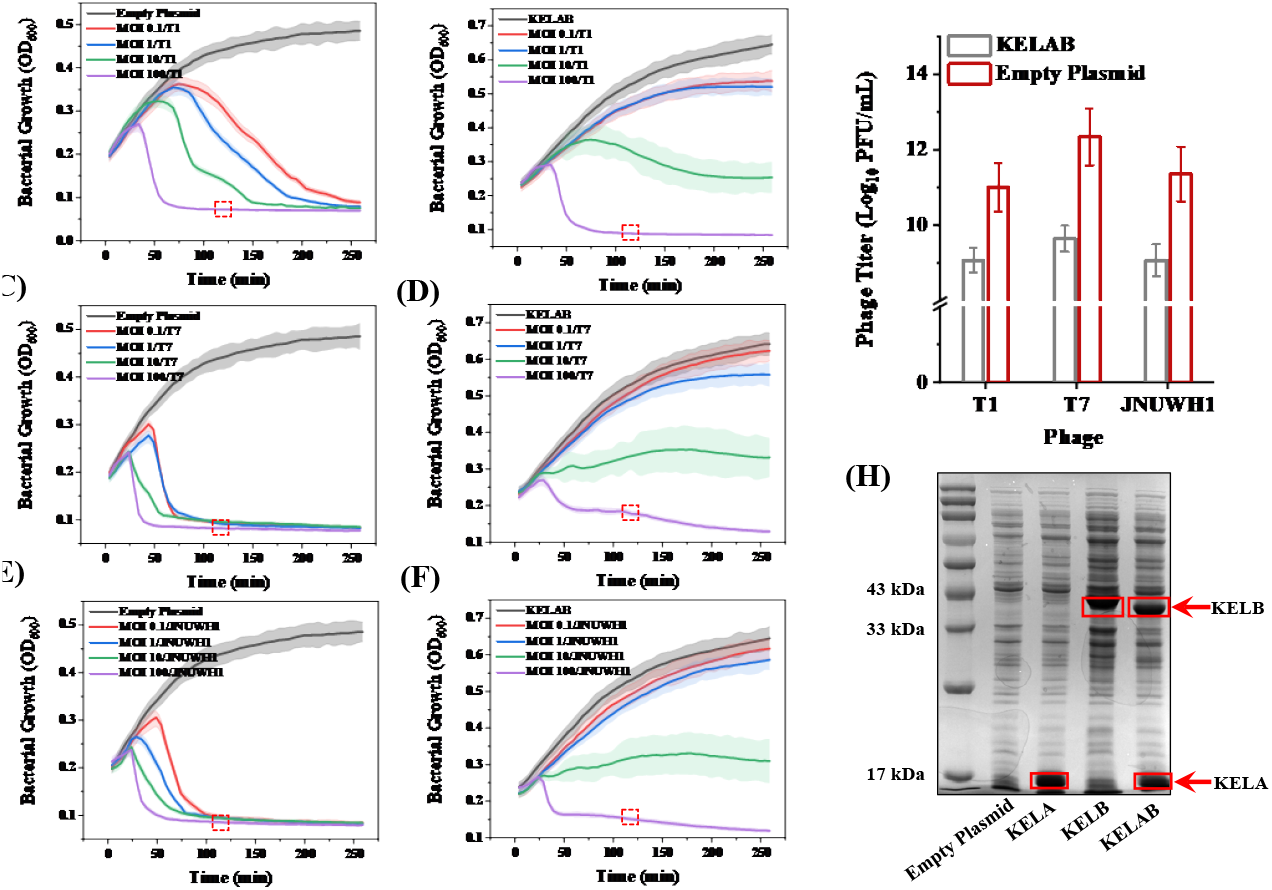
KELAB confers protection against bacteriophage infection. (A-F) Growth curves of *E. coli* expressing KELAB in response to bacteriophage infections at different multiplicities of infection (MOIs) with phages T1, T7, and JNUWH1. The curves represent the average of two independent experiments, each performed in triplicate. Error bars indicate the standard deviation of the mean. (G) Magnification of the area marked with a dashed rectangle in (A-F). To detect the release of phage progeny by dead cells with or without the KELShedu defense system, the lysates of dead cells with phage infection at MOI 100 (120 min post-infection for T1, T7 and JNUWH1) from the above assays were collected. Then Phage titer in lysate was determined. Bar graphs represent the mean ± standard deviation (SD) from three independent experiments. (H) SDS-PAGE analysis of protein expression in *E. coli* BL21(DE3) harboring either the empty pET-28a plasmid, KELA, KELB and the KELAB gene cassette.

We assessed the expression of KELAB proteins, and SDS-PAGE analysis confirmed the concurrent expression of KELA and KELB. Although we hypothesize that KELA might function as a transcriptional repressor similar to BrxR or CapW, it did not inhibit the expression of KELB (Fig. 2H). Furthermore, the presence of KELAB did not impose significant growth constraints on *E. coli* in the absence of phage infection. These findings suggest that the KELShedu system represents a novel abortive infection-based anti-phage mechanism with phage recognition capability.

### KELB forms a complex with KELA in vitro and in vivo

Multicomponent bacterial defense systems like Gabija and BREX typically have components that cooperate at the molecular scale(*5, 6*). To elucidate the mechanisms by which KELA and KELB confer protection to *E. coli*, we first verified the in vivo protein-protein interaction between these two proteins encoded by the KELAB operon. When expressing KELAB operon, KELA, with an N-terminal His-tag, was purified using Ni-NTA, co-purifying KELB due to their interaction (Fig. 3A). MALDI-TOF analysis confirmed the co-purified band as KELB, indicating an interaction with KELA. Subsequently, we validated the in vitro interaction through pull-down assays. SDS-PAGE and MALDI-TOF results confirmed that KELB interacts with KELA in vitro as well (Fig. 3B). These in vivo and in vitro interaction assays suggest that KELA and KELB form a complex that functions to protect *E. coli* against phage infection.

**Figure 3.**
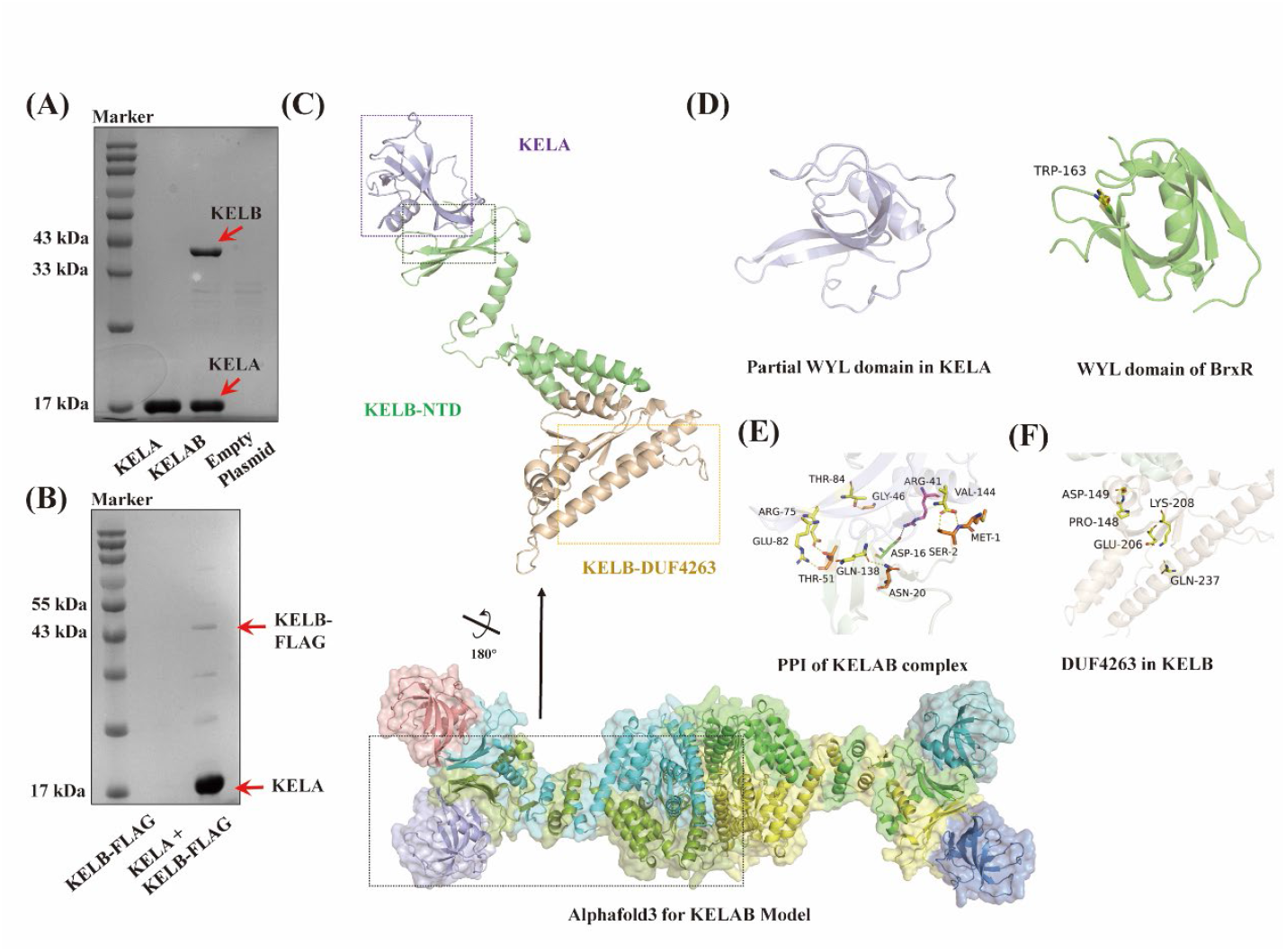
KELA and KELB form a complex in vivo and in vitro. (A) In vivo verification of protein-protein interactions between KELA and KELB. When His-tagged KELA is purified from *E. coli* harboring the KELAB gene cassette, its co-precipitates with non-His-tagged KELB, demonstrating the interaction between these proteins. (B) In vitro verification of the protein-protein interaction between KELA and KELB. Pull-down assays demonstrate that KELB interacts with KELA in vitro, confirming their interaction outside the cellular environment. (C) AlphaFold server modeling of the KELAB complex. In heterooctamers, we assign different polymers to different colors. In a separate KELAB, The model highlights KELA in purple, the N-terminal domain of KELB in green, and the C-terminal DUF4263 domain of KELB in yellow. (D) KELA is similarity to the WYL domain, a Sm-type SH3 β-barrel fold. However, KELA lacks the tryptophan residue essential for a canonical WYL domain of BrxR (PDB ID 7t8k). (E) Protein complex interaction (PPI) analysis of KELAB complex. (F) KELB’s DUF4263 domain, associated with the PD-(D/E)XK phosphodiesterase superfamily.

To further investigate the interaction dynamics of KELAB proteins, we used the AlphaFold Server(*21*) to model heterooctamer of KELA and KELB according to the result of Agilent Bio SEC. The structural analysis revealed that the KELAB complex exhibits an overall candy-like structure, with KELB subunits arranged in pairs. The protein interaction interface is mediated by a C-terminal α-helical bundle in KELB, while KELA binds to the N-terminal region of KELB on the external surface of the complex (Fig. 3C). KELA comprises a β-sheet structure formed by four β-strands. The results of structural alignments using the DALI server(*22*) showed that KELA had Z-scores of 11.8 and 11.0 with CapW (PDB ID 7tb5) and BrxR (PDB ID 7t8k), respectively. Notably, studies indicate that CapW (*23*)and BrxR(*12*) are members of the universal defense signaling protein family, playing roles in sensing phage gene invasion in CBASS and BREX systems, respectively. Dali server structural comparisons indicated that KELA is similarity to the WYL domain, a Sm-type SH3 β-barrel fold(*12*). However, KELA lacks the tryptophan residue essential for a canonical WYL domain, raising questions about its typical WYL functionality (Fig. 3D). In contrast, KELB showed Z-scores of 7.6 and 7.4 with Card1 nuclease (PDB ID 6wxx) and endonuclease (PDB ID 5gke), respectively. KELB’s N-terminal domain (NTD) features a β-sheet structure (residues 1-55) that mediates binding to KELA (Fig. 3E), while residues 55-77 form a hinge leading to the C-terminal DUF4263 domain. Structural modeling suggests that KELA may mediate recognition of small molecules like DNA, while KELB’s DUF4263 domain, associated with the PD-(D/E)XK phosphodiesterase superfamily (Fig. 3F). Given its DUF4263 domain and association with the PD-(D/E)XK phosphodiesterase superfamily, known to be involved in replication, restriction, DNA repair, and tRNA-intron splicing(*24*), we speculate that KELB possesses nuclease activity.

This detailed structural and interaction analysis underscores the potential functional roles of KELA and KELB in the KELShedu system, highlighting their collaborative mechanism in providing phage resistance. Further experimental validation and functional assays are warranted to fully elucidate the mechanistic underpinnings and biological significance of this novel abortive infection system.

### Functional Divergence of KELA and KELB in DNA Binding and Hydrolysis

To elucidate the functions of KELA and KELB, we investigated their DNA-binding capabilities. EMSA results demonstrated that KELA altered the migration pattern of linearized pET-28a plasmid DNA on polyacrylamide gel, indicating that KELA binds to linear double-stranded DNA (dsDNA, Fig. 4A). In contrast, KELB did not show significant binding to linear dsDNA, possibly due to its nuclease activity, which might hydrolyze the DNA, preventing it from being visualized on the gel.

**Figure 4.**
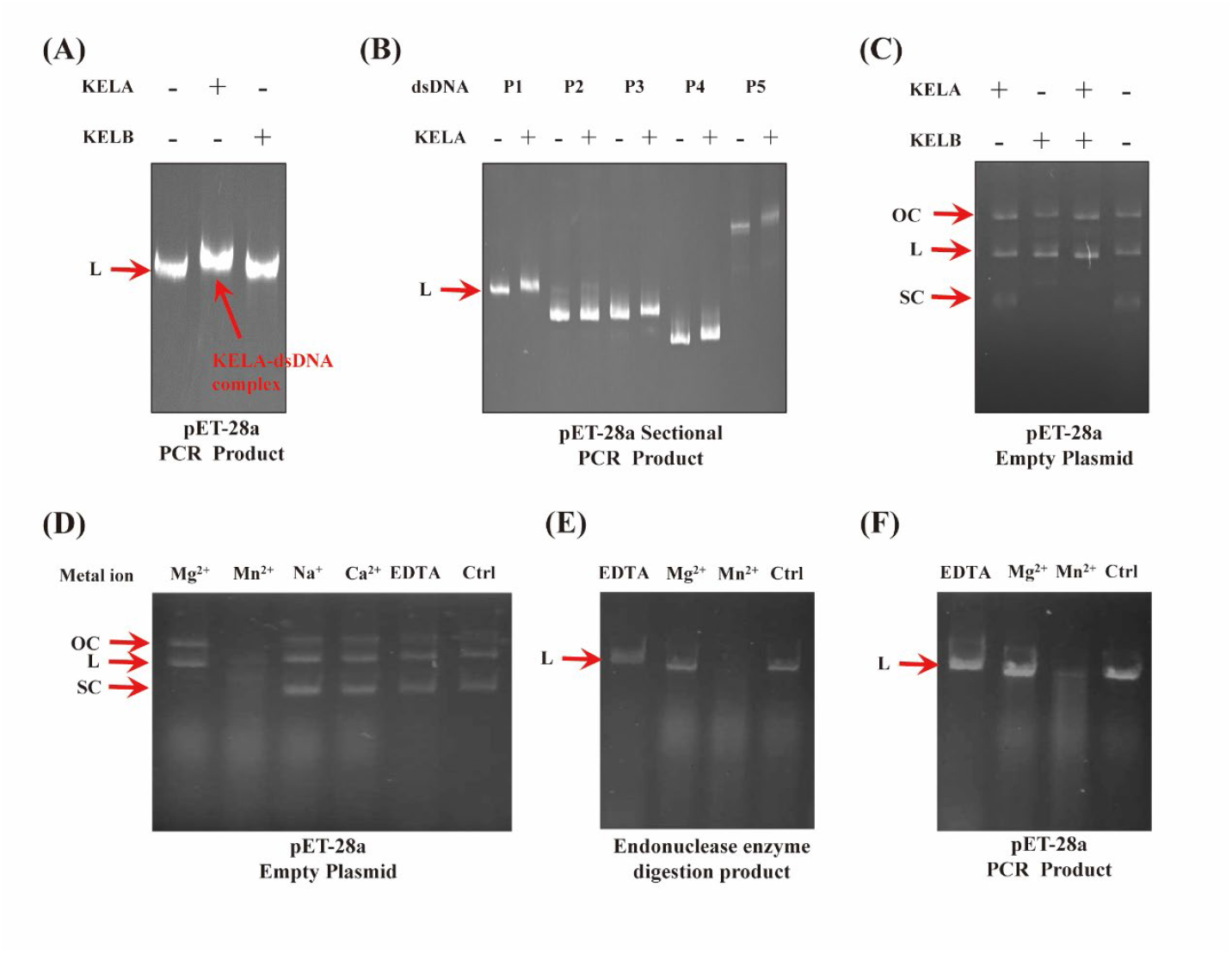
DNA Binding and Nuclease Activity of KELA and KELB. (A) EMSA Analysis of KELA-DNA Binding: EMSA demonstrating KELA’s binding to dsDNA. Incubation of KELA and KELB with PCR-linearized pET28a plasmid, followed by EMSA analysis. (B) Non-specific DNA Recognition by KELA: EMSA showing KELA’s binding to five PCR-generated fragments of pET28a plasmid (P1-P5), indicating non-specific recognition of dsDNA. (C) Nuclease Activity of KELB in Presence of Mg^2+^: Validation of nuclease activity of KELA, KELB, and KELAB in the presence of magnesium ions. (D) Metal Ion Dependency of KELB Nuclease Activity: Assessment of KELB’s nuclease activity in the presence of various metal ions, highlighting its metal-dependent nature. (E) Nuclease Activity on Modified pET28a Fragments: Evaluation of KELB’s nuclease activity on restriction enzyme-digested pET-28a plasmid fragments in the presence of Mg^2+^ and Mn^2+^ ions, demonstrating activity on biologically modified DNA. (F) Nuclease Activity on Unmodified PCR Fragments: Validation of KELB’s nuclease activity on unmodified, PCR-linearized pET-28a fragments in the presence of Mg^2+^ and Mn^2+^ ions, confirming its activity on unmodified DNA. Images are representative of three replicates.

To further explore KELA’s dsDNA recognition specificity, we divided the pET-28a plasmid into five segments (P1-P5) of 1000-2000 base pairs each through PCR and assessed KELA’s binding to these segments. EMSA results showed that KELA bound to all five segments, suggesting non-specific recognition of dsDNA (Fig. 4B). Additionally, EMSA with varying concentrations of purified KELA and linearized pET-28a fragments demonstrated a progressive decrease in DNA migration with increasing KELA amounts (Fig. S3A), confirming non-specific dsDNA binding by KELA (Fig. S3B). This non-specific binding capacity of KELA may indicate a broader regulatory or structural role within the cell, potentially aiding the KELAB complex in executing various DNA-related functions.

To further verify whether KELB with the PD-(D/E)XK phosphodiesterase domain has nuclease cleavage activity, we investigated the DNA hydrolysis properties using a plasmid digestion assay in the presence of magnesium ions, which are commonly used in nuclease buffer solutions(*6*). The results showed that KELA did not exhibit DNA hydrolysis activity in the presence of magnesium ions, whereas KELB demonstrated cleavage activity on supercoiled plasmid DNA (SC, Fig. 4C). Moreover, the addition of KELA did not affect the cleavage activity of KELB. Next, we tested the cleavage effect of KELB on linearized pET-28a DNA obtained by PCR. Agarose gel electrophoresis results indicated that KELB did not exhibit nuclease activity on linearized dsDNA in the presence of magnesium ions (Fig. S4A).

Further investigation into the effect of metal ions on KELB activity revealed that KELB cleaved supercoiled, nicked, and linearized forms of pET-28a DNA in the presence of manganese ions (Fig. 4D). However, no nuclease activity was observed in the presence of Na^+^, Ca^2+^ or EDTA (Fig. 4D). To explore KELB’s cleavage pattern in the presence of magnesium and manganese ions, we conducted time-course cleavage reactions (Fig. S4B). No shorter cleavage products were detected, indicating that KELB is not an exonuclease and does not have specific cleavage sites.

Based on previous research, nucleases in DISARM(*25*) and BREX(*4, 26*) systems recognize specific restriction-modification sites. To determine if KELB has a similar mechanism, we performed digestion assays on pET-28a plasmid DNA extracted from active *E. coli* BL21(DE3) using *Bam*H I and *Hin*d III restriction endonucleases. This method preserves the modifications made by *E. coli* BL21(DE3) in a biologically active state. Digestion assays in the presence of manganese ions showed that KELB exhibited nuclease activity on both double-restriction enzyme-digested (Fig. 4E) and PCR-linearized dsDNA (Fig. 4F). These results suggest that KELB is a metal-dependent nuclease without sequence-specific recognition sites. Additionally, to further ascertain that the nuclease activity of KELB is mediated through the DUF4263 domain, we mutated its conserved Q237 residue(*27*) to determine the critical site for Shedu antiphage activity. The KELABQ237A mutant lost its antiphage activity (Fig. S5), and nuclease assays revealed that the mutant protein no longer exhibited nuclease activity (Fig. S6).

This comprehensive analysis highlights the metal-ion dependency and non-specific DNA hydrolysis properties of KELB. However, further research is required to elucidate how the KELShedu system defends against bacteriophages.

### The Impact of KELShedu system on Cellular DNA Levels

Based on previous experiments, we observed that expressing KELAB proteins does not inherently activate the abortive infection response of the KELShedu system. This raises the question: how does the KELShedu system trigger Abi activity in *E. coli* carrying the KELShedu system upon phage invasion? To explore this, we stained the cell membrane and intracellular

DNA with fluorescent dyes and used laser confocal microscopy to observe the effects of T7 phage infection on bacterial cells expressing KELAB. Prior to phage invasion, the bacterial genomic DNA was localized in a concentrated, fixed region within the cell. However, within a very short time after phage invasion (less than 15 minutes), the genomic DNA in KELAB-expressing bacteria became evenly dispersed throughout the cell. This dispersion may indicate structural disruption of the genomic DNA. As time progressed, the DNA in cells expressing the KELShedu system was gradually degraded (Fig. 5A)(*28*). These observations suggest that the KELShedu system induces significant changes in the cellular DNA upon sensing phage invasion, leading to DNA degradation as part of the Abi mechanism.

**Figure 5.**
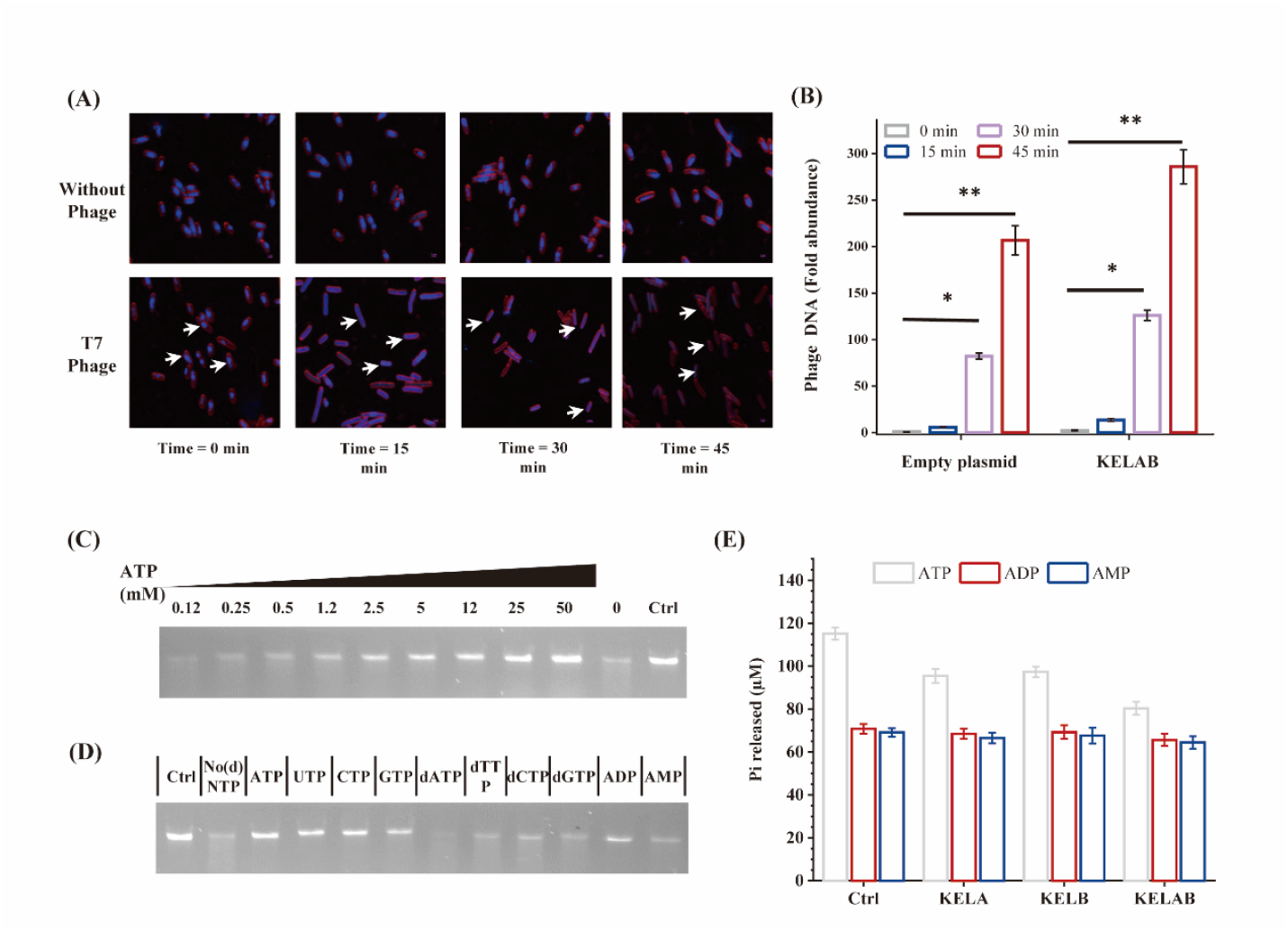
KELshedu Degrades Intracellular DNA When ATP Levels Decline. (A) *E. coli* BL21(DE3) cells harboring the KELShedu system were infected with phage T7 at a multiplicity of infection (MOI) of 2, and their cellular responses were observed at different time points post-infection using laser confocal microscopy. DAPI staining was employed to visualize DNA, depicted in indigo, while FM4-64 staining highlighted the cell membrane, shown in magenta. Imaging conditions and parameters were maintained consistently across all samples. The white arrows represent the degradation of the host DNA. The images presented are representative of three independent replicates. (B) The relative abundance of phage DNA and host DNA at various time points post-infection was quantified using qPCR. (C) Inhibition of KELAB nuclease activity by varying concentrations of ATP. (D) Effects of physiological concentrations (5 mM) of NTP, dNTP, ADP, and AMP on KELAB nuclease activity. (E) Assessment of ATPase, activity in KELA, KELB, and the KELAB complex, the concentration of ATP, ADP and AMP was 5 mM. Images are representative of three replicates. Bar graphs represent the mean ± standard deviation (SD) from three independent experiments. Statistical significance was determined using Student’s t-test, with *p < 0.05 and **p < 0.005.

To further investigate the dynamics of phage and host DNA within the cell, we extracted total DNA from *E. coli* carrying either an empty vector or the KELShedu system at different time points following phage invasion. We then used qPCR to analyze the bacterial 16s rRNA gene and the phage-specific tail fiber protein gene, tracking changes in phage and host DNA levels over time. The results showed that, compared to cells lacking KELAB, those expressing KELAB did not exhibit a significant difference in phage DNA levels (Fig. 5B), indicating that the phage replication cycle continued unimpeded after invasion. This indicates that the KELShedu system does not initiate DNA degradation immediately upon phage DNA injection. Instead, it non-specifically degrades all intracellular DNA, including both genomic DNA and phage DNA, during the replication phase of phage DNA. Gel electrophoresis analysis of DNA integrity at different time points post-invasion further confirmed that the KELShedu system degrades DNA following phage attack.

To further investigate the de-repression mechanism of KELShedu following phage invasion, we observed that physiological concentrations of ATP (2.5–15 mM) inhibit the DNA cleavage activity of the KELAB complex (Fig. 5C). This phenomenon suggests that KELShedu may share a similar activation mechanism with systems like Gabija(*6, 29*) and Septu(*30*). We then examined the dsDNA cleavage activity of KELAB in the presence of UTP, CTP, GTP, dNTPs, ADP, and AMP. The results showed that physiological concentrations of NTPs and ADP inhibit KELAB’s cleavage activity, while dNTPs have no effect. This implies that the KELShedu anti-phage defense system is activated by sensing a drop in intracellular NTP levels. Further investigation into the impact of ATP on KELA-dsDNA binding and KELB’s dsDNA cleavage revealed that ATP does not affect KELA’s binding to dsDNA (Extended Figure 7A), whereas ATP inhibits KELB’s cleavage activity in a manner similar to the KELAB complex (Extended Fig. 7B).

These in vivo and in vitro findings demonstrate that the KELShedu anti-phage system de-represses its dsDNA cleavage activity by sensing a decrease in intracellular NTP levels. Additionally, Although KELA and KELB do not have ATPase related domains from the domain retrieval results, we still tested the ATPase activity of KELA and KELB. We confirmed that neither KELA, KELB, nor the KELAB complex degrades ATP (Fig. 6E). This highlights the role of NTPs in our model, acting as inhibitory factors that suppress KELAB activity under normal physiological conditions. Upon phage invasion, the drop in intracellular NTP levels due to phage replication and metabolic demands activates the nucleic acid cleavage activity of KELShedu, ultimately leading to abortive infection.

**Figure 6.**
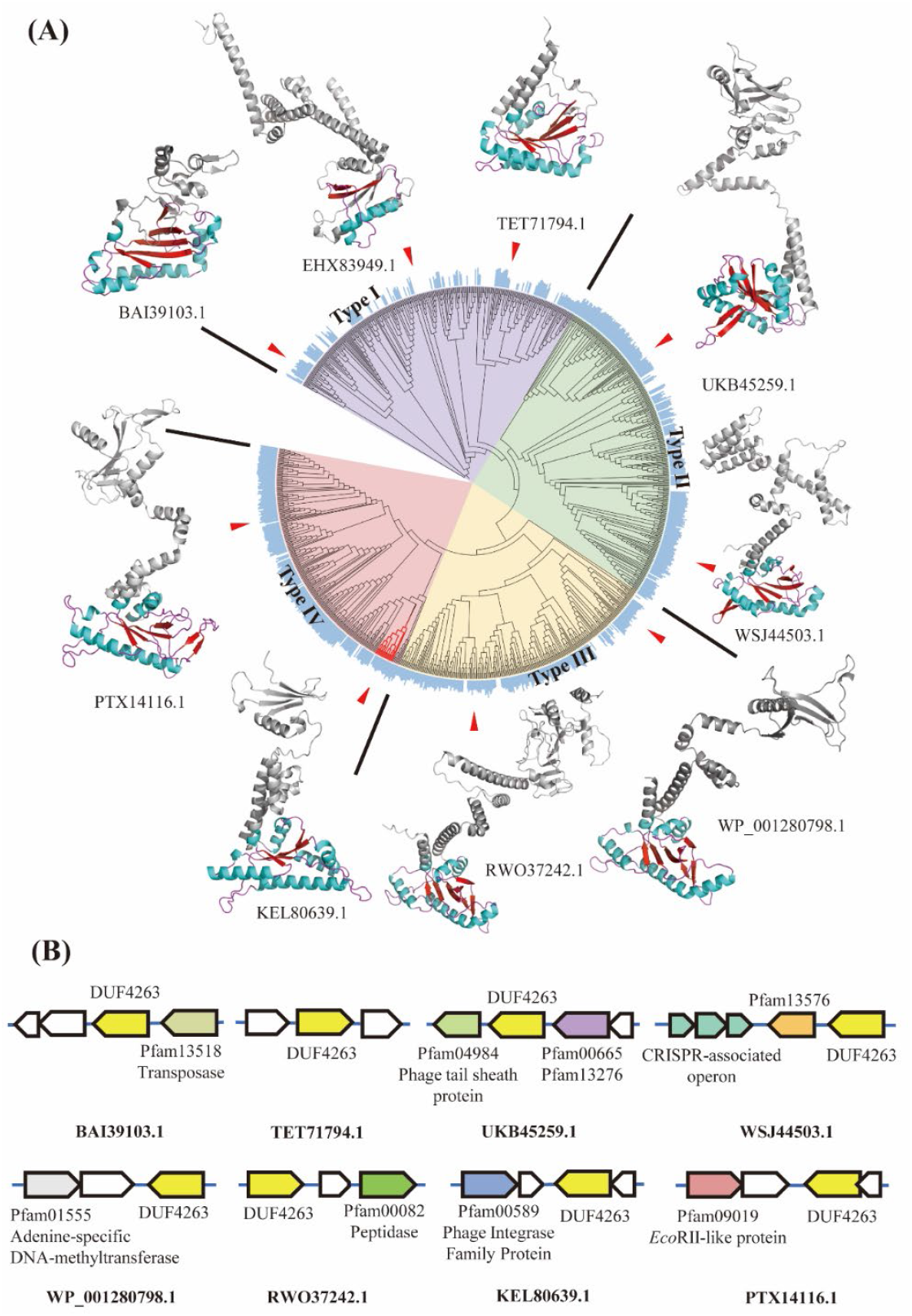
Phylogenetic Analysis and Genomic Localization of the Shedu System. (A) Phylogenetic tree analysis of proteins containing the DUF4263 domain, categorizing the Shedu system into four distinct subtypes (Type I-IV) based on evolutionary branching. Within the protein structures, the colored regions represent the DUF4263 domain, with α-helices shown in cyan and β-sheets in red. The likelihood of being classified as a 3.1.21.X nuclease is depicted as bar graphs in the outer ring of the phylogenetic tree. (B) Genomic localization of Shedu proteins, with different domain-containing proteins represented by distinct colors in the diagram. Yellow indicates proteins containing the DUF4263 domain, while white denotes hypothetical proteins with uncharacterized domains.

### Shedu Belongs to a Superfamily of Defense Systems with Four Major Subtypes

The results from our experiments, alongside previous studies, suggest that the Shedu system represents a phage defense mechanism with significant mechanistic complexity. To gain a deeper understanding of this system and its evolutionary relatives, we constructed a phylogenetic tree using proteins containing the Shedu system’s core domain, DUF4263 (Data S1), as shown in Fig. 6A. The phylogenetic relationships depicted in this tree indicate a clear progression in the complexity of the Shedu system, from small, single-component proteins to larger, multi-component systems.

This evolutionary trend is vividly illustrated in the structural representations surrounding the tree, with colored regions denoting the position of the DUF4263 domain within each protein. During extensive evolutionary processes, we categorized the Shedu system into four major subtypes. The purple portion represents the ancestral Type I classification of Shedu proteins, exemplified by BAI39103.1 and TET71794.1, which display streamlined and minimalistic Shedu proteins with only 2-3 scattered α-helices or β-sheets, aside from the essential DUF4263 domain. Notably, EHX83949.1, which was previously classified as a Shedu protein by Doron et al., did not exhibit significant anti-phage activity in prior experiments. Structural comparison reveals that, unlike typical DUF4263 domains, it contains only partial α-helix and β-sheet aggregations in its tertiary structure. Moreover, EFPNET predictions for nuclease activity (EC 3.1.21.X) demonstrate a widespread loss of nuclease function in Type I, highlighting the functional limitations of ancestral proteins.

As evolution progressed, the Shedu system diverged into two subtypes: Type II (green) and Type III (yellow). These subtypes likely arose through horizontal gene transfer and other evolutionary events, leading ancestral Shedu proteins to incorporate additional domains, forming larger and more complex proteins. In both Type II and Type III, the DUF4263 domain predominantly localizes to the C-terminus of Shedu proteins, while the N-terminal domains often cannot be identified as known conserved domains. Analysis through AlphaFold server modeling reveals, for instance, that in UKB45259.1 and WSJ44503.1, the N-terminal domain is connected to the DUF4263 domain at the C-terminus via an α-helix-loop hinge. In the broader clustering context, Type III and the subsequent Type IV (discussed below) share similar structures, which may be indicative of flexible interactions between the N- and C-terminal domains or potentially related to the oligomeric states of Shedu proteins(*15*). In WSJ44503.1, the N-terminus is composed of an antiparallel 4-α-helix bundle, while UKB45259.1 features an α-helix-β-sheet-α-helix sandwich structure. A similar arrangement is observed in RWO37242.1 of Type III, albeit with more disordered loop regions. WP_001280798.1, on the other hand, has a 5-β-sheet topped by an α-helix at its N-terminus.

In the classification of Shedu systems as Type IV, more refined N-terminal structures are observed. Both PTX14116.1 and KEL80639.1 are identified as candidates within this group, appearing as dual-component systems (Fig. 6B). While the upstream operon functions of PTX14116.1 remain unknown, the evolutionary trajectory of the Shedu system is quite clear.

However, it is important to note that not all Type IV proteins are organized as multi-operon systems.

Phylogenetic analysis suggests that early Shedu systems likely originated from small, single-domain proteins that gradually evolved into larger, more complex proteins. Ultimately, these proteins appear to have developed into multi-component systems, potentially enhancing their functionality in bacterial phage defense and emphasizing the critical role of the N-terminal domain throughout evolution.

Fig. 6B provides additional context by mapping the genomic locations of these Shedu proteins, which are predominantly situated near other defense-related genes. This genomic arrangement suggests that Shedu systems are typically found within defense islands enriched with diverse defense mechanisms, potentially facilitating coordinated responses to phage attacks.

Additionally, the dispersion of Shedu systems from the same Enterobacteriaceae origins across the phylogenetic tree without clear evolutionary relationships supports the notion of widespread horizontal gene transfer. This horizontal movement may be a key factor in the dissemination of Shedu systems across different bacterial species, contributing to their evolutionary success and functional diversification.

## Discussion

Our investigation provides insights into the KELShedu system’s unique mechanism of phage resistance, distinguishing it as a nucleotide-responsive defense system within the Shedu family. KELShedu employs an abortive infection strategy, where both genomic and phage DNA are degraded non-specifically in response to phage-triggered intracellular changes, effectively disrupting phage replication cycles and protecting the bacterial population. This response is tightly linked to nucleotide depletion, as physiological concentrations of ATP and other NTPs inhibit the DNA cleavage activity of the KELAB complex. Only upon a reduction in intracellular NTP levels does the KELShedu system activate its DNA degradation function. (Fig. 7)

**Figure 7.**
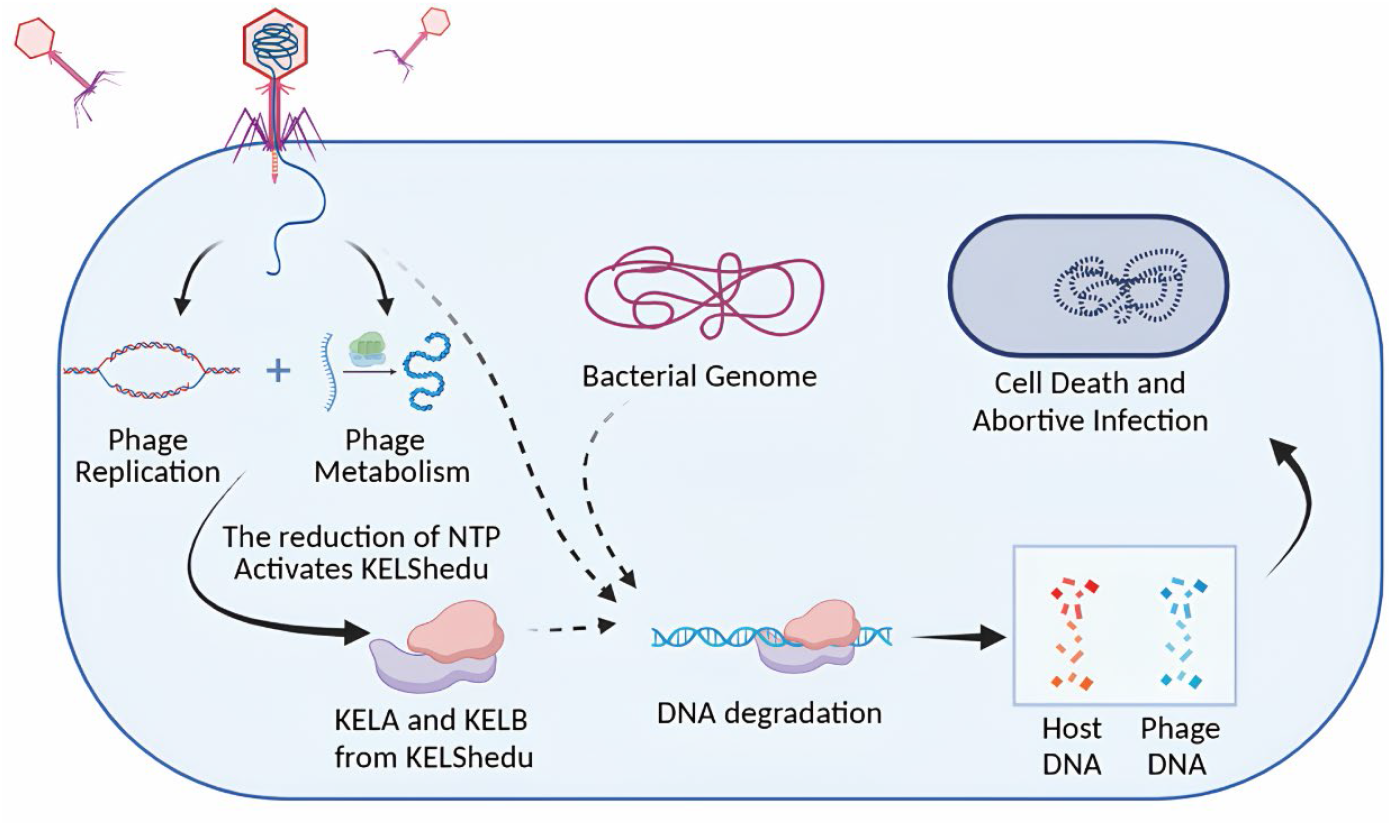
Construction of KELshedu Anti-phage System Model. Following phage invasion, intracellular NTP levels are significantly depleted due to the phage’s replication and metabolic activities. This reduction in NTP concentration effectively lifts the inhibition on KELAB nuclease activity. Consequently, KELAB induces the degradation of intracellular DNA, leading to cell death. Through this abortive infection mechanism, the KELShedu system protects the bacterial population as a whole by halting phage propagation.

In comparing KELShedu to other known anti-phage systems, such as Gabija(*6, 7, 27, 29*), Septu(*30*), and CBASS(*8, 14*), we observe distinct regulatory nuances that set KELShedu apart. The Gabija and Septu systems, for instance, also couple their nuclease activity to cellular nucleotide conditions. Gabija, like KELShedu, responds to reduced nucleotide levels, but unlike the straightforward ATP inhibition in KELShedu, Gabija incorporates a multi-component assembly and specialized recognition domains that distinguish phage invasion through secondary signals. Septu, on the other hand, is activated in response to phage-induced stress markers rather than nucleotide depletion directly, utilizing a more complex sensor-effector arrangement to detect phage presence and trigger cellular defense responses. In contrast, CBASS relies on cyclic oligonucleotide signaling pathways to activate its defense effectors, a mechanism fundamentally different from KELShedu’s direct nucleotide-sensing strategy.

Moreover, our study highlights the need for a broader exploration of Shedu system diversity. Recent studies by Yajie Gu (*15*), and colleagues on Shedu systems from *B. cereus* demonstrate variations in anti-phage mechanisms that could indicate divergent evolutionary strategies within the Shedu family. Such comparative studies are crucial to understanding the versatility of Shedu-mediated defenses across different bacterial species. By examining how other Shedu systems function across diverse microbial hosts, we may uncover additional regulatory and functional innovations that enhance bacterial immunity against phages.

In summary, while the current study provides foundational insights into the KELShedu system’s operation, future efforts focused on structural and comparative analyses are essential. Detailed structural studies will clarify the precise mechanisms by which KELAB responds to phage-induced nucleotide depletion, while comparative studies with other Shedu systems will deepen our understanding of Shedu evolution and functional diversity. Together, these insights will support the development of KELShedu-based applications in microbial engineering and synthetic biology, advancing bacterial resilience against phage threats

## Materials and Methods

### Construction, Expression and Purification

The KELAB gene cluster was synthesized by Sangon Biotech (Shanghai) Co., Ltd. To preserve the ribosome binding site (RBS) of KELB, codon optimization was applied except for the 36 bp upstream of the KELB start codon. Post-synthesis, the genes were amplified via PCR to obtain the KELA and KELB fragments and inserted into the *Bam*H I and *Hin*d III sites of the pET-28a plasmid using ClonExpress Ultra One Step Cloning Kit. The KELB-FLAG fusion, created by adding a FLAG tag to the C-terminus of KELB through PCR, was inserted into the *Nco* I and *Hin*d III sites. The constructed plasmids were introduced into *E. coli* BL21(DE3) using the CaCl_2_ method. Mutant strains were generated by amplifying the wild-type genes with primers carrying the mutation and assembling the fragments through ClonExpress Ultra One Step Cloning Kit. High-fidelity PCR enzymes and ClonExpress Ultra One Step Cloning Kit were sourced from Vazyme International LLC., *Bam*H I and *Hin*d III were sourced from Takara Biomedical Technology (Beijing) Co., Ltd., while the plasmids and *E. coli* BL21(DE3) were lab stock.

The constructed strains were cultured in 10 mL of LB medium containing 50 mg/mL kanamycin at 37°C for 8 hours. Subsequently, they were transferred at a 5% inoculation ratio to 50 mL of LB medium and grown at 37°C until an OD_600_ of 0.5. Protein expression was induced with 0.5 mM IPTG at 16°C for 12 hours with shaking at 180 rpm. After induction, cells were harvested by centrifugation at 8000 rpm for 10 minutes, resuspended in PBS, and centrifuged again to remove impurities. The cell pellet was ultrasonically disrupted (250 W, 1s pulse, 3s pause, 15 minutes), and the lysate was centrifuged at 12000 rpm for 30 minutes to remove cell debris. The supernatant was filtered through a 0.22 µm membrane and purified using Ni-NTA agarose. The Ni-NTA column (Dongfulong Qianchun Biotechnology (Shanghai) Co., LTD) was pre-equilibrated with 10 column volumes of elution buffer M0 (20 mM Tris-HCl, 500 mM NaCl, pH 7.4) and washed with 15 column volumes of M0 containing 20 mM and 80 mM imidazole.

The protein was eluted with M0 containing 250 mM imidazole. The collected eluate was dialyzed using a Millipore Amicon Ultra-15 (10,000 MWCO) to remove most of the imidazole. All reagents are purchased from Macklin Inc.

### Phage Plaque Assays

To purify the phage using the double-layer agar method(*31*), 100 µL of the phage suspension was diluted to approximately 10^3^ pfu/mL and mixed with 300 µL of *E. coli* culture (OD_600_ = 0.6-0.8) in 5 mL of semi-solid LB medium containing 0.5% agar. The mixture was then poured onto solid LB medium containing 1% agar and incubated overnight at 37°C.

Subsequently, individual purified phage plaques were carefully picked and introduced into *E. coli* culture (OD_600_ = 0.6-0.8) for amplification. The lysate was centrifuged at 8000 rpm for 10 minutes and filtered through a 0.22 µm membrane, then stored in SM buffer at 4°C to ensure optimal stability for subsequent experiments. The lysate was serially diluted from 10^-1^ to 10^-8^ in tenfold steps, and the phage titer was determined using the double-layer agar method.

For phage plaque assays, 300 µL of *E. coli* culture (OD_600_ = 0.8-1.0) was mixed with 5 mL of semi-solid LB medium containing 0.1 mM IPTG and 50 mg/mL kanamycin, and then poured onto solid LB medium. After solidification, the plates were incubated at 25°C for 1.5 hours to induce protein expression. A 1 µL aliquot of the phage dilution was spotted onto the agar overlay. The plates were then incubated inverted at 25°C for 4 - 8 hours.

### Measurement of Bacterial Growth Curve

The bacterial strains were inoculated into 10 mL of LB medium containing 50 mg/mL kanamycin and cultured at 37°C for 6 hours. Subsequently, 50 µL of the cell culture was added to a 96-well microtiter plate containing LB medium supplemented with 0.5 mM IPTG and 50 mg/mL kanamycin. Phages were added at the desired multiplicity of infection (MOI). The plates were incubated with shaking at 25°C, and the OD_600_ was measured every 3 minutes for a duration of 6 hours using a Biotek Epoch2 microplate reader (Agilent Technologies, Inc.). This experiment was performed in triplicate to ensure reproducibility, with error bars indicating the standard error of the mean.

### Adsorption Rate of the Bacteriophage

Based on previous research(*31*), to calculate the adsorption rate of the bacteriophage, the *E. coli* strain used in this study was cultured until it reached the early logarithmic growth phase (OD_600_ = 0.6-0.8). The bacteriophage was then added to the bacterial culture at a MOI of 1, followed by incubation at 37°C. As a control(S0), the same procedure was performed using LB medium without bacteria. Samples were collected after 15 minutes and centrifuged to separate the supernatant, which was subsequently diluted to an appropriate concentration. The diluted supernatant was then mixed with *E. coli* BL21(DE3) using the double-layer agar plate method and incubated at 37°C for 6 hours. The phage titer(S) was determined by counting the number of individual plaques formed with error bars indicating the standard error of the mean. The adsorption rate of the bacteriophage was calculated using the formula:

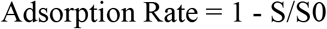

### Protein Interaction Assays

To validate the in vivo interaction between KELA and KELB, we utilized a purification approach analogous to co-immunoprecipitation (co-IP) using Ni-NTA chromatography on strains co-expressing the KELAB operon(*6*), where KELA was expressed with a His-tag, while KELB was untagged (methodology as described previously). The control group consisted of strains expressing only His-tagged KELA, which were purified similarly.

For the in vitro interaction validation of KELA and KELB, we employed a His-pulldown assay. KELA was expressed and purified using Ni-NTA chromatography. The columns were washed with 15 column volumes of M_0_ buffer containing 20 mM and 80 mM imidazole, respectively. The filtered KELB-FLAG cell lysate was then incubated with the Ni-NTA agarose column for 15 minutes. After incubation, the columns were washed again with 15 column volumes of M0 buffer containing 20 mM and 80 mM imidazole, respectively, and the bound proteins were eluted with M_0_ buffer containing 250 mM imidazole. For the control, cell lysate from the pET-28a empty vector strain was processed similarly to replace the KELA lysate. The collected eluates were analyzed by SDS-PAGE to verify the presence of the proteins.

Protein identification was performed using a Bruker Daltonics MALDI-TOF mass spectrometer. The corresponding bands from the SDS-PAGE gel were excised, destained, digested with trypsin, and then subjected to mass spectrometric analysis. Protein identification was conducted using the Matrix Science database.

### Electrophoretic Mobility Shift Assay

Electrophoretic Mobility Shift Assay (EMSA) was performed using a 6.5% native polyacrylamide gel(*32*). The electrophoresis buffer was 0.5x TBE. Prior to sample loading, the gel was pre-electrophoresed at 90 V on ice for 90 minutes. The samples were prepared by mixing 2x binding solution, 5 nM of protein, 100 ng of DNA, and double-distilled water to a final volume of 20 µL. The mixture was incubated at room temperature for 10 minutes before loading onto the gel. Electrophoresis was conducted at 120 V on ice for 1.5-2 hours. After electrophoresis, the gel was carefully transferred to 0.5x TBE buffer containing 2 µL of Goldview Nucleic Acid Gel Stain (Yisheng Biotechnology (Shanghai) Co., LTD) to visualize the DNA bands.

### Nuclease Assays

To evaluate the nuclease activity of KELA and KELB, DNA cleavage assays were conducted at 37°C. The reaction buffer consisted of 10 mM Tris-HCl, 250 mM NaCl, and pH 7.4, with the addition of 5 mM MgCl_2_, MnCl_2_, CaCl_2_, or EDTA to determine optimal reaction conditions. After selecting the optimal conditions, 150 ng of DNA substrate and 3 nM protein were incubated in the reaction buffer. The reactions were performed at 37°C for 30 minutes and terminated by adding a stop solution containing 25 mM EDTA and 2.5 mg/mL proteinase K and performed at 37°C for 10 minutes. Samples were analyzed by native agarose gel electrophoresis or 6% non-denaturing polyacrylamide gel. Quantitative bar charts represent the average of three independent experiments.

### Laser Confocal Microscopy

Laser confocal microscopy was performed as previously described. Briefly, *E. coli* BL21(DE3) containing either the KELAB gene cassette were grown overnight at 37°C and subsequently inoculated at 5% into fresh media until reaching an OD600 of 0.3-0.4. The cultures were then infected with phage T7 at a multiplicity of infection (MOI) of 2. Samples were collected at 0-, 10-, 20-, and 30-minutes post-infection. The cells were stained with 4′,6-diamidino-2-phenylindole (DAPI; 4 mg/ml) for DNA imaging and FM4-64 (2 mg/ml) for membrane imaging. For each condition, more than 10 fields from each of two independent experiments were manually scored, resulting in data from over 1,000 cells for each condition.

### Quantitative real-time PCR

*E. coli* BL21(DE3) cells carrying either the KELAB gene cassette or an empty vector were infected with phage T7 at an MOI of 0.5, with samples collected at 0-, 15-, 30-, and 45-minutes post-infection. Immediately following collection, samples were heated at 100°C for 10 minutes to lyse cells and release DNA. Equal volumes of each sample were used for qPCR analysis. qPCR was performed using ChamQ SYBR qPCR Master Mix (Q311, Vazyme) and the StepOnePlus Real-Time PCR System. DNA abundance at 0 minutes post-infection was normalized to the control group. The ΔΔCt method was employed to calculate differences between the control (empty vector, 0 min) and sample (ΔCt), with results expressed as 2^(-ΔΔCt)^. The relative DNA copy number at each time point post-infection was normalized against the control (empty vector). Each experiment was conducted in triplicate across at least three independent trials.

### Genomic Data and Identification of the Shedu System

We screened a subset of proteins containing the DUF4263 domain from the NCBI database, filtering these proteins from a pool of over 2 billion sequences based on their domain features. Subsequently, we constructed a phylogenetic tree of these proteins using the Neighbor-Joining method implemented in the MEGA 11.0 software. To ensure the robustness and reliability of the phylogenetic relationships, we further evaluated the tree using Bootstrap Replications. This phylogenetic analysis allowed us to thoroughly investigate the evolutionary relationships among these proteins, revealing potential divergence pathways and functional linkages that could provide critical theoretical support for subsequent biological research.

Following the construction of the phylogenetic tree, we employed the enzyme function prediction model EFPNET, which was developed using deep learning techniques, to predict the enzymatic functions of the amino acid sequences. Specifically, we built a dataset comprising over 220,000 sequences with experimentally validated enzymatic functions from the Swiss-Prot database, categorizing them into more than 4,900 functional classes based on EC numbers. We used the ESM and Prot-T5 protein language models to perform multi-view embedding of these enzyme sequences, integrating a part-of-speech enhancement module through a self-attention mechanism. The model utilized Swish as the activation function and Focal-Loss as the loss function to optimize prediction outcomes. The final model outputs were transformed into probability distributions via the Softmax function, with output values as real numbers between 0 and 1, ensuring that the sum of probabilities across all categories equaled one (unpublished).

We then applied EFPNET to test the Shedu data from our phylogenetic tree. The results indicated that the majority of sequences were predicted to be nucleases, specifically EC number 3.1.21.X. Following validation through wet-lab experiments, we confirmed the accuracy of these predictions. The predicted probabilities of sequences being nucleases (EC 3.1.21.X) were represented as outer-circle data on the phylogenetic tree diagram, with non-nuclease sequences assigned a value of zero.

## Acknowledgments

This work was supported by grants from the National Natural Science Foundation of China (No. 32071470), the Postgraduate Research & Practice Innovation Program of Jiangsu Province (KYCX23_2501), Jiangsu Program for Frontier Technology R&D (BF2024012) and the Project Funded by the Priority Academic Program Development of Jiangsu Higher Education Institutions, Top-notch Academic Programs Project of Jiangsu Higher Education Institutions.

## Author contributions

Conceptualization: JY, XZ, Hengwei Zhang

Methodology: ZZ, Hanwei Zhang, HZhang, HW, DZ

Investigation: HZhang, ZZ

Visualization: HZhang, JY

Supervision: XP, JY, ZR, WZ

Writing—original draft: HZhang

Writing—review & editing: HZhang, JY, ZR, XZ

## Data and materials availability

All data are available in the main text or the supplementary materials.

## Supplementary Materials

**This PDF file includes:**

Figs. S1 to S7

Tables S1

**Other Supplementary Materials for this manuscript include the following:**

Data S1

## References

1. G. Gasiunas, T. Sinkunas, V. Siksnys, Molecular mechanisms of CRISPR-mediated microbial immunity. Cellular and Molecular Life Sciences 71, 449–465 (2013).

2. C. Rouillon, N. Schneberger, H. Chi, K. Blumenstock, S. Da Vela, K. Ackermann, J. Moecking, M. F. Peter, W. Boenigk, R. Seifert, B. E. Bode, J. L. Schmid-Burgk, D. Svergun, M. Geyer, M. F. White, G. Hagelueken, Antiviral signalling by a cyclic nucleotide activated CRISPR protease. Nature 614, 168–174 (2022).

3. S. Song, Y. Guo, J.-S. Kim, X. Wang, T. K. Wood, Phages Mediate Bacterial Self-Recognition. Cell Reports 27, 737–749.e734 (2019).

4. J. Gordeeva, N. Morozova, N. Sierro, A. Isaev, T. Sinkunas, K. Tsvetkova, M. Matlashov, L. Truncaitė, R. D. Morgan, N. V. Ivanov, V. Siksnys, L. Zeng, K. Severinov, BREX system ofEscherichia colidistinguishes self from non-self by methylation of a specific DNA site. Nucleic Acids Research 47, 253–265 (2019).

5. T. Goldfarb, H. Sberro, E. Weinstock, O. Cohen, S. Doron, Y. Charpak‐Amikam, S. Afik, G. Ofir, R. Sorek, BREX is a novel phage resistance system widespread in microbial genomes. The EMBO Journal 34, 169–183 (2014).

6. R. Cheng, F. Huang, X. Lu, Y. Yan, B. Yu, X. Wang, B. Zhu, Prokaryotic Gabija complex senses and executes nucleotide depletion and DNA cleavage for antiviral defense. Cell Host & Microbe 31, 1331–1344.e1335 (2023).

7. S. P. Antine, A. G. Johnson, S. E. Mooney, A. Leavitt, M. L. Mayer, E. Yirmiya, G. Amitai, R. Sorek, P. J. Kranzusch, Structural basis of Gabija anti-phage defence and viral immune evasion. Nature 625, 360–365 (2023).

8. B. Lowey, A. T. Whiteley, A. F. A. Keszei, B. R. Morehouse, I. T. Mathews, S. P. Antine, V. J. Cabrera, D. Kashin, P. Niemann, M. Jain, F. Schwede, J. J. Mekalanos, S. Shao, A. S. Y. Lee, P. J. Kranzusch, CBASS Immunity Uses CARF-Related Effectors to Sense 3′– 5′- and 2′–5′-Linked Cyclic Oligonucleotide Signals and Protect Bacteria from Phage Infection. Cell 182, 38–49.e17 (2020).

9. W. E. Robertson, L. F. H. Funke, D. de la Torre, J. Fredens, T. S. Elliott, M. Spinck, Y. Christova, D. Cervettini, F. L. Böge, K. C. Liu, S. Buse, S. Maslen, G. P. C. Salmond, J. W. Chin, Sense codon reassignment enables viral resistance and encoded polymer synthesis. Science 372, 1057-+ (2021).

10. Y. P. Liu, L. Dai, J. H. Dong, C. Chen, J. G. Zhu, V. B. Rao, P. Tao, Covalent Modifications of the Bacteriophage Genome Confer a Degree of Resistance to Bacterial CRISPR Systems. Journal of Virology 94, (2020).

11. D. M. Picton, Y. A. Luyten, R. D. Morgan, A. Nelson, Darren L. Smith, D. T. F. Dryden, J. C. D. Hinton, Tim R. Blower, The phage defence island of a multidrug resistant plasmid uses both BREX and type IV restriction for complementary protection from viruses. Nucleic Acids Research 49, 11257–11273 (2021).

12. Y. A. Luyten, D. E. Hausman, J. C. Young, L. A. Doyle, K. M. Higashi, N. C. Ubilla-Rodriguez, A. R. Lambert, C. S. Arroyo, K. J. Forsberg, R. D. Morgan, B. L. Stoddard, B. K. Kaiser, Identification and characterization of the WYL BrxR protein and its gene as separable regulatory elements of a BREX phage restriction system. Nucleic Acids Research 50, 5171–5190 (2022).

13. D. M. Picton, J. D. Harling-Lee, S. J. Duffner, S. C. Went, R. D. Morgan, J. C. D. Hinton, T. R. Blower, A widespread family of WYL-domain transcriptional regulators co-localizes with diverse phage defence systems and islands. Nucleic Acids Research 50, 5191–5207 (2022).

14. B. Duncan-Lowey, N. K. McNamara-Bordewick, N. Tal, R. Sorek, P. J. Kranzusch, Effector-mediated membrane disruption controls cell death in CBASS antiphage defense. Molecular Cell 81, 5039–5051.e5035 (2021).

15. Y. Gu, H. Li, A. Deep, E. Enustun, D. Zhang, K. D. Corbett, Bacterial Shedu immune nucleases share a common enzymatic core regulated by diverse sensor domains. (2023).

16. S. Doron, S. Melamed, G. Ofir, A. Leavitt, A. Lopatina, M. Keren, G. Amitai, R. Sorek, Systematic discovery of antiphage defense systems in the microbial pangenome. Science 359, (2018).

17. X. Zou, X. H. Xiao, Z. R. Mo, Y. S. Ge, X. Jiang, R. L. Huang, M. X. Li, Z. X. Deng, S. Chen, L. R. Wang, S. Y. Lee, Systematic strategies for developing phage resistant strains. Nature Communications 13, (2022).

18. S. Wang, E. Sun, Y. Liu, B. Yin, X. Zhang, M. Li, Q. Huang, C. Tan, P. Qian, V. B. Rao, P. Tao, The complex roles of genomic DNA modifications of bacterioph. (2022).

19. A. Lopatina, N. Tal, R. Sorek, Abortive Infection: Bacterial Suicide as an Antiviral Immune Strategy. Annual Review of Virology 7, 371–384 (2020).

20. A. Millman, A. Bernheim, A. Stokar-Avihail, T. Fedorenko, M. Voichek, A. Leavitt, Y. Oppenheimer-Shaanan, R. Sorek, Bacterial Retrons Function In Anti-Phage Defense. Cell 183, 1551–1561.e1512 (2020).

21. J. Abramson, J. Adler, J. Dunger, R. Evans, T. Green, A. Pritzel, O. Ronneberger, L. Willmore, A. J. Ballard, J. Bambrick, S. W. Bodenstein, D. A. Evans, C. C. Hung, M. O’Neill, D. Reiman, K. Tunyasuvunakool, Z. Wu, A. Zemgulyte, E. Arvaniti, C. Beattie, O. Bertolli, A. Bridgland, A. Cherepanov, M. Congreve, A. I. Cowen-Rivers, A. Cowie, M. Figurnov, F. B. Fuchs, H. Gladman, R. Jain, Y. A. Khan, C. M. R. Low, K. Perlin, A. Potapenko, P. Savy, S. Singh, A. Stecula, A. Thillaisundaram, C. Tong, S. Yakneen, E. D. Zhong, M. Zielinski, A. Zídek, V. Bapst, P. Kohli, M. Jaderberg, D. Hassabis, J. M. Jumper, Accurate structure prediction of biomolecular interactions with AlphaFold 3. Nature, (2024).

22. L. Holm, A. Laiho, P. Törönen, M. Salgado, DALI shines a light on remote homologs: One hundred discoveries. Protein Sci 32, (2023).

23. Q. S. Liang, S. T. Richey, S. N. Ur, Q. Z. Ye, R. K. Lau, K. D. Corbett, Structure and activity of a bacterial defense-associated 3′-5′ exonuclease. Protein Sci 31, (2022).

24. K. Steczkiewicz, A. Muszewska, L. Knizewski, L. Rychlewski, K. Ginalski, Sequence, structure and functional diversity of PD-(D/E)XK phosphodiesterase superfamily. Nucleic Acids Research 40, 7016–7045 (2012).

25. G. Ofir, S. Melamed, H. Sberro, Z. Mukamel, S. Silverman, G. Yaakov, S. Doron, R. Sorek, DISARM is a widespread bacterial defence system with broad anti-phage activities. Nature Microbiology 3, 90–98 (2017).

26. K. Bhushan, BacteRiophage EXclusion (BREX): A novel anti‐phage mechanism in the arsenal of bacterial defense system. Journal of Cellular Physiology 233, 771–773 (2017).

27. S. S. Wang, E. Sun, Y. P. Liu, B. Q. Yin, X. Q. Zhang, M. L. Li, Q. Huang, C. Tan, P. Qian, V. B. Rao, P. Tao, Landscape of New Nuclease-Containing Antiphage Systems in Escherichia coli and the Counterdefense Roles of Bacteriophage T4 Genome Modifications. Journal of Virology 97, (2023).

28. R. K. Lau, Q. Ye, E. A. Birkholz, K. R. Berg, L. Patel, I. T. Mathews, J. D. Watrous, K. Ego, A. T. Whiteley, B. Lowey, J. J. Mekalanos, P. J. Kranzusch, M. Jain, J. Pogliano, K. D. Corbett, Structure and Mechanism of a Cyclic Trinucleotide-Activated Bacterial Endonuclease Mediating Bacteriophage Immunity. Molecular Cell 77, 723–733.e726 (2020).

29. R. Cheng, F. Huang, H. Wu, X. Lu, Y. Yan, B. Yu, X. Wang, B. Zhu, A nucleotide-sensing endonuclease from the Gabija bacterial defense system. Nucleic Acids Research 49, 5216–5229 (2021).

30. Y. Y. Li, Z. F. Shen, M. Y. Zhang, X. Y. Yang, S. P. Cleary, J. L. Xie, I. A. Marathe, M. Kostelic, J. Greenwald, A. D. Rish, V. H. Wysocki, C. Chen, Q. Chen, T. M. Fu, Y. M. Yu, PtuA and PtuB assemble into an inflammasome-like oligomer for anti-phage defense. Nature Structural & Molecular Biology 31, (2024).

31. H. Zhang, J. You, X. Pan, Y. Hu, Z. Zhang, X. Zhang, W. Zhang, Z. Rao, Genomic and biological insights of bacteriophages JNUWH1 and JNUWD in the arms race against bacterial resistance. Frontiers in Microbiology 15, (2024).

32. L. M. Hellman, M. G. Fried, Electrophoretic mobility shift assay (EMSA) for detecting protein-nucleic acid interactions. Nat Protoc 2, 1849–1861 (2007).

