## Supplementary material for "The Two-Component Nuclease-Active KELShedu System Confers Broad Anti-Phage Activity via Abortive Infection": SM

Hengwei Zhang *et al.*

**This PDF file includes:**

Figs. S1 to S7
Tables S1

**Other Supplementary Materials for this manuscript include the following:**

Data S1

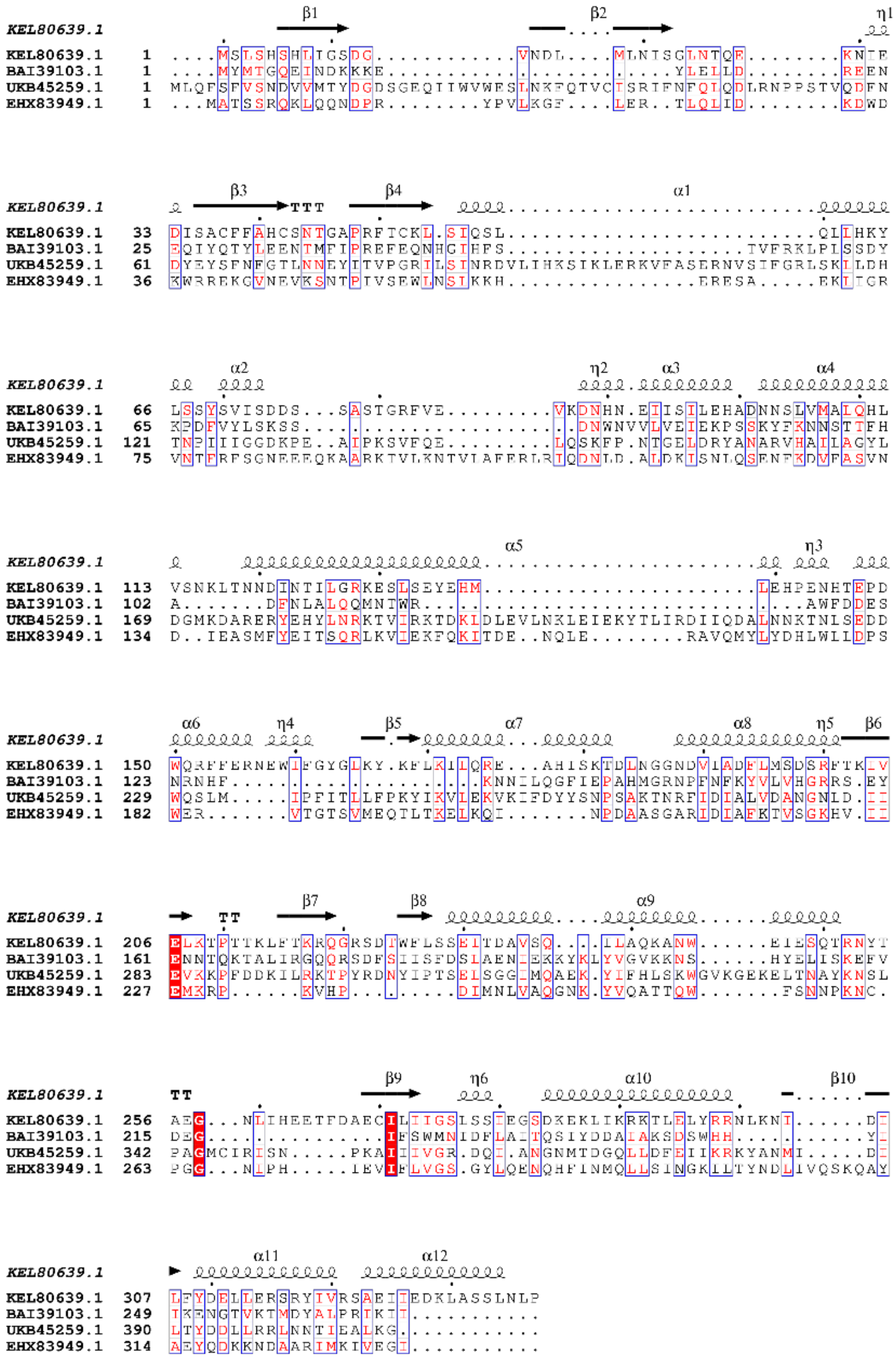

**Fig. S1. Selected Proteins with The DUF4263 Domain from Various Enterobacteriaceae Sources.** We identified four Shedu proteins: KEL80639.1, BAI39103.1, UKB45259.1, EHX83949.1 and the sequence differences were also aligned.

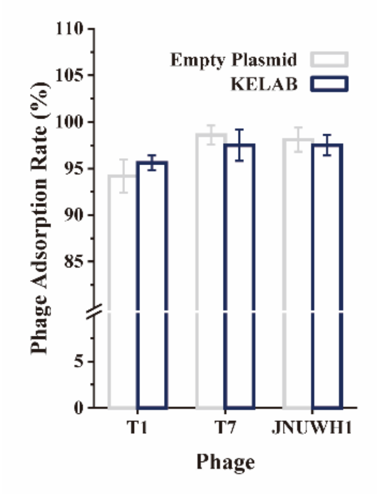


**Fig. S2. Determination of Phage Adsorption Rate:** Adsorption rates of bacteriophages T1, T7, and JNUWH1 to *E. coli* BL21(DE3) strains harboring the KELshedu system versus those containing an empty plasmid. The data represent the mean of three independent replicates, with error bars indicating the standard deviation of the mean.

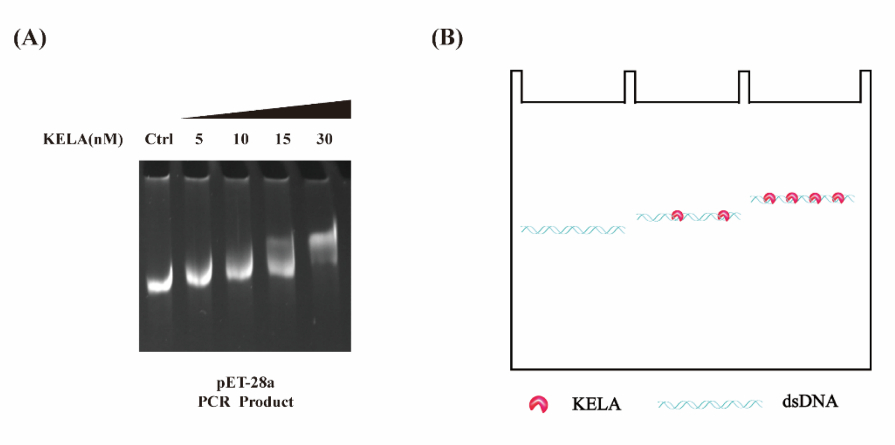


**Fig. S3. EMSA analysis of KELA-DNA binding:** (A) EMSA showed different concentrations of KELA binding to dsDNA. Ctrl for the control group without the addition of KELA protein. Each sample contained 100 ng pET-28a PCR product (B) Different concentrations of KELA lead to changes in DNA band mobility, indicating the amount of DNA binding to the KELA protein

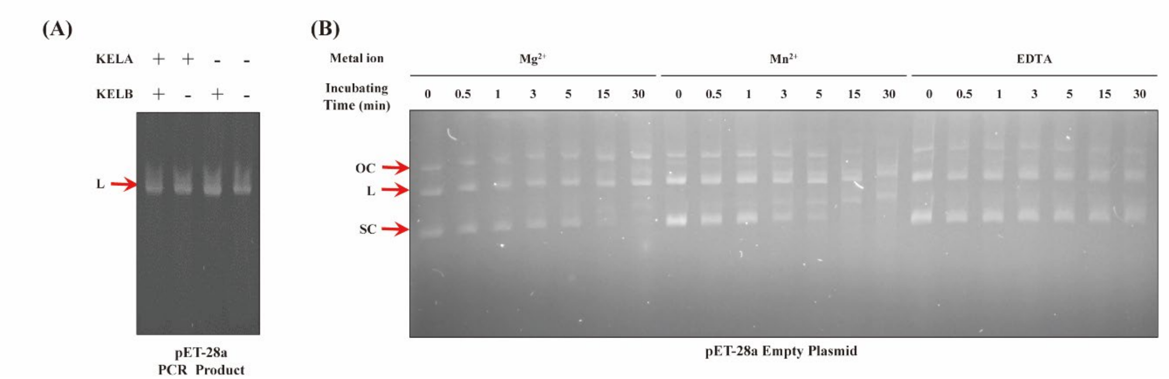


**Fig. S4. Nuclease Activity of KELA and KELB.** (A) The cleavage effect of KELB on linearized pET-28a DNA obtained by PCR in the presence of magnesium ions. (B) The nuclease effect of KELB with time in the presence of magnesium and manganese ions

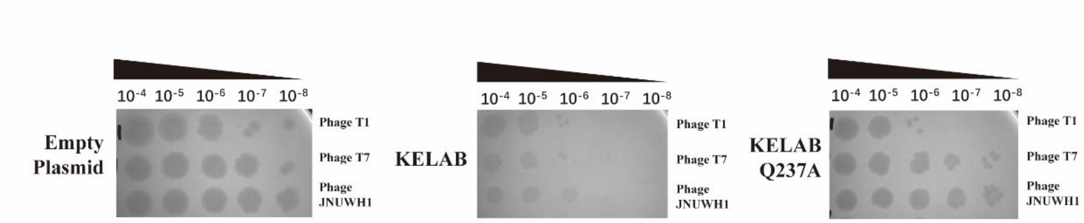


**Fig. S5. Mutated KELShedu System Lost Bacteriophage Resistance to *E. coli*.**
Phage resistance in *E. coli* BL21(DE3) expressing different ORFs of the KELAB cassette. Strains containing empty vector, KELShedu system, and mutated KELShedu system were tested. Displayed are the results of plaque assays with tenfold serial dilutions of phages T1, T7, and JNUWH1. The images are representative of three independent replicates.

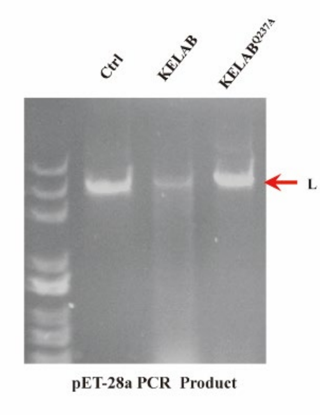


**Fig. S6. Nuclease Assays.** Result revealed that the Mutant KELB No Longer Exhibited Nuclease Activity.

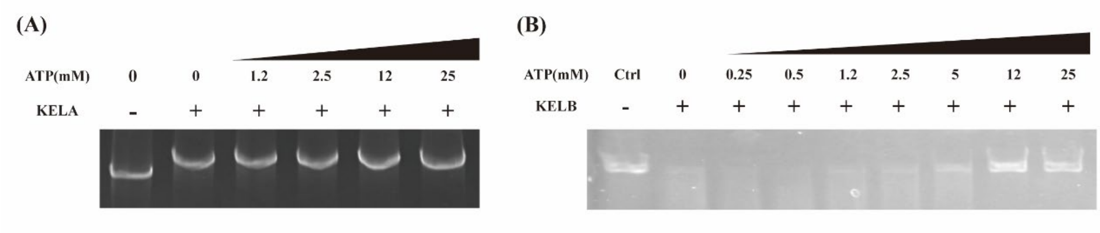


**Fig. S7. The effect of ATP on KELABshedu system.** (A) EMSA analysis of the effect of varying ATP concentrations on KELA-dsDNA binding. (B) Effect of varying ATP concentrations on KELB-mediated dsDNA cleavage.

**Tables S1**

**Primers used for construction of plasmids, site-directed mutagenesis and RT-qPCR.**The primer name, sequence, and application are shown below.

| Primer Name | Sequence | Application |
|---|---|---|
| 28a-KEL-F | cagcaaatgggtcgcggatccatgagtctacaattattgagggaatcagtacaac | Construction of plasmids |
| 28a-KEL-R | tcgagtgcggccgcaagcttttatggcagattcaaagagctcgc | Construction of plasmids |
| 28a-KB-flag-R-1 | tcatcatcgtccttatagtccttatcatcatcatccttgtaatccttgtcgtcgtcgtcc<br>ttgtagtctggcagattcaaagagctcgc | Construction of plasmids |
| 28a-KB-flag-R-2 | gagtgcggccgcaagcttttacttgtcatcatcgtccttatagtccttatcatcatcat | Construction of plasmids |
| 28a-KA- -R | gctcgagtgcggccgcaagcttttatacctttgctatcacattttgataaaaatcggac | Construction of plasmids |
| 28a-1-F | gatgcgtccggcgtagagg | Fragment of dsDNA |
| 28a-5-R | attatttctagaggggaattgttatccgctc | Fragment of dsDNA |
| 28a-2-F | tgcaatttattcatatcaggattatcaataccatatttttgaaaa | Fragment of dsDNA |
| 28a-1-R | tatggaactgcctcggtgagt | Fragment of dsDNA |
| 28a-3-F | cagcagagcgcagataccaaat | Fragment of dsDNA |
| 28a-2-R | agttcttgaagtggtggcctaact | Fragment of dsDNA |
| 28a-4-F | cctgtttggtcactgatgcctc | Fragment of dsDNA |

| Primer Name | Sequence | Application |
| --- | --- | --- |
| 28a-3-R | atcggtatcattacccccatgaacagaaa | Fragment of dsDNA |
| 28a-5-F | gagacgggcaacagctgat | Fragment of dsDNA |
| 28a-4-R | ggcaaaccagcgtggacc | Fragment of dsDNA |
| T7-rt-F | CTTTGGAGGCGTGGGGTTAT | RT-qPCR |
| T7-rt-R | GTTCCACCACTCCATTCCGT | RT-qPCR |
| E.coli-16s-F | TAGTAGGTGGGGTAACGGCT | RT-qPCR |
| E.coli-16s-R | GGACCGTGTCTCAGTTCCAG | RT-qPCR |
